# Dose-response relationships of LSD-induced subjective experiences in humans

**DOI:** 10.1101/2022.11.05.515283

**Authors:** Tim Hirschfeld, Johanna Prugger, Tomislav Majić, Timo T. Schmidt

## Abstract

Lysergic acid diethylamide (LSD) is a potent classic serotonergic psychedelic, which facilitates a variety of altered states of consciousness. Here we present the first meta-analysis establishing dose-response relationship estimates of the altered experience induced by LSD. Data extracted from articles identified by a systematic literature review following PRISMA guidelines were obtained from the Altered States Database. The psychometric data comprised ratings of subjective effects from standardized and validated questionnaires: the Altered States of Consciousness Rating Scale (5D-ASC, 11-ASC) and the Mystical Experience Questionnaire (MEQ30). We performed meta-regression analyses using restricted cubic splines for data from studies with LSD doses of up to 200 μg base. Most scales revealed a sigmoid-like increase of effects with a plateauing at around 100 μg. The most strongly modulated factors referred to changes in perception and illusory imagination, followed by positively experienced ego-dissolution, while only small effects were found for *Anxiety* and *Dread of Ego Dissolution*. The partly observed considerable variability of effects points to the importance of non-pharmacological effects on subjective experiences. The established dose-response relationships may be used as general references for future experimental and clinical research on LSD to relate observed with expected subjective effects and to elucidate phenomenological differences between psychedelics.

## Introduction

D-lysergic acid diethylamide (LSD) is the prototype of classic serotonergic psychedelics, a group of substances which unfold their psychoactive properties predominantly via the serotonin 2A (5-HT_2A_) receptor [1]. Psychedelics embrace structurally heterogenous subgroups like phenethylamines (e.g., mescaline) and tryptamines (e.g., psilocybin, *N,N*-DMT) [2], as well as substances from the ergoline subgroup (e.g., LSD) which have been characterized as “rigidified tryptamines” [3]. The term ‘psychedelics’ is also used in a broader sense, including non-serotonergic drugs like Ketamine, PCP or MDMA. The term ‘psychedelic experience’ is used in an even broader sense, not limited to the effects induced by specific substances, instead referring to a group of psychological effects. However, there is no clear definition on the exact set of consciousness alterations that define a psychedelic experience. Here, we will refer to classic serotonergic psychedelics and the effects they induce when using the term ‘psychedelics’ or ‘psychedelic experience’. Several studies suggest that qualitatively, LSD might not be differentiated from other psychedelics with regard to the induced psychologic effects [4–6]. On the other hand, anecdotal reports mention differences in subjective experiences regarding different substances [7], and LSD somewhat differs from pharmacodynamical profiles of other 5-HT_2A_ agonists, including a broader variety of receptor targets [3].

After its initial synthesis in 1938, LSD’s psychedelic properties have accidentally been discovered in 1943 by the Swiss pharmacologist Albert Hofmann [8]. Ever since, LSD has been the most extensively investigated psychedelic from the 1950s to the 1970s, with more than 1,000 scientific papers published in the context of basic science, as well as in clinical research as a therapeutic tool [9–11]. Most intensively studied indications included, among others, alcoholism [12] and existential distress in life-threatening physical illness [13]. After a hiatus of more than 20 years, during which regulatory hurdles prevented research on psychedelics, research eventually resumed in the 1990s. Also, in recreational underground use LSD is by far the most frequently used psychedelic worldwide [14], and dosages of psychedelics are often compared to LSD equivalents by users.

LSD has recently been re-evaluated for the treatment of different mental health conditions, like anxiety and depression in patients with [15,16] and without life-threatening illness [15]. Psychological effects of psychedelics underlie specific temporal dynamics [9], including (acute) psychedelic experiences, subacute effects, and long-term (enduring) effects [17]. There is some evidence that the quality and intensity of acute psychedelic effects might predict therapeutic outcome [18]. Thus, the classification and description of acute psychedelic experiences appear to be of high importance when it comes to optimizing treatment interventions regarding efficacy and safety. In order to determine the optimal dosing ranges of LSD for future clinical studies, first and most importantly the influence of the LSD dose on the nature and intensity of acute subjective effects needs characterization. Over the last decades some gold standards for the assessment of altered states phenomena have been established in terms of several well-validated questionnaires for an retrospective assessment of altered experiences [9,19–21]. Such standardized assessment allows meta-analytic comparisons, as recently presented to establish dose-response relationships for altered experiences induced by psilocybin [20].

The data to establish dose-response relationships for LSD is limited by the availability of studies that cohere to current research standards, as LSD has only recently returned to basic and clinical research. The range of LSD doses employed in current research is on average markedly below the doses administered in studies from the 1950-70s. At that time, high doses mostly ranging between 400 and 600 μg were applied, especially in the treatment of alcohol use disorder, whereas in recent studies doses of 200 μg have not been exceeded, neither in healthy subjects [22], nor in clinical samples [15,16,23]. Regarding the lower end of the dose spectrum, applications of so-called microdosing are gaining increasing interest both in recreational and scientific settings. It is thought that such low doses, which do not induce subjectively identifiable drug effects, can positively affect cognitive processes and improve mental health indicators [24–27]. The exact dose range of LSD microdosing is still debated; a recent literature review suggests a “plausible microdose range” for LSD of 6-20 μg [24]. Only few studies directly tested the dose-dependency of LSD’s effects [22,26,28]. With the application of within-subject designs, these studies provide well-controlled experimental conditions, while they do not allow to assess study-specific impacts on participants experiences. Such non-pharmacological effects appear, however, particularly pronounced for psychedelic substances, and are generally referred to as set and setting-effects [29,30]. Cross-study comparisons are essential to obtain data-based estimates of the variability of subjective LSD experiences across research sites and experimental protocols, as no corresponding meta-analysis is available to date.

With the present meta-analysis, we aim to obtain estimates of the relationship between LSD doses and the quality of psychedelic experiences. The data stem from the Altered States Database (ASDB, http://alteredstatesdb.org, [31]), which is a regularly updated database with questionnaire data extracted from articles identified by a systematic literature review, adhering to the Preferred Reporting Items for Systematic Reviews and Meta-Analyses (PRISMA) 2020 Statement Guidelines [32]. Available data was found for the Altered States of Consciousness Rating Scale (5D-ASC, 11-ASC) and the Mystical Experience Questionnaire (MEQ30) for a dose range up to 200 μg.

## Methods

### Included data

Psychometric questionnaire data on the subjective experience of LSD were included in this meta-analysis. The data has been retrieved from the ASDB repository on Open Science Framework (OSF; https://osf.io/8mbru, version ASDB_v.2022-12-31), which contains data from MEDLINE-listed studies on altered states of consciousness published from 1975-01-01 until 2022-12-31. The ASDB stems from a systematic literature review according to PRISMA standards [32] as described in Prugger et al. (2022) [33] and Peters et al. (2023) [34]. Datasets investigating the subjective effects of LSD were retrieved from the ASDB. The PRISMA flowchart showing the process of item identification and screening can be found in the Supplementary Material.

Data were excluded if experimental conditions comprised applications of combinations of substances [28,35,36] such as pre-treatments with ketanserin, if the LSD dose was unclear [37], or if data were about recreational LSD usage [38]. From studies reporting multiple questionnaire applications at different time points during the same day of the experimental session [39,40], only the final and complete questionnaire application was included describing the overall experience. Also, after consultation with the authors, the LSD doses in a series of reports [22,35,36,39,41] were adjusted as suggested in [28,42], due to administration of capsules containing an unstable LSD formulation leading to dispersion of lower than presumed LSD doses. Studies reporting on administration of both LSD base and LSD tartrate have been included. LSD tartrate doses (from studies [43–45]) were converted to the LSD base equivalent by multiplying by 0.685 [46] (See Table 1).

**Table 1.** Summary of studies included in the meta-regression analyses. Table 1 displays the studies included in the meta-regression analysis, with sample size of study participants, study description (study design, additional conditions, physiological assessment and additional measurements), questionnaire used to report data, LSD administration method, and LSD base dose (*converted from tartrate doses). Several studies contain multiple observations (e.g., from repeated measurements). Note: LSD: lysergic acid diethylamide; ECG: electrocardiogram; EEG: electroencephalogram; fMRI: functional magnetic resonance imaging; MDMA: 3,4-methylenedioxy-methamphetamine; MEG: Magnetoencephalography; PO: per oral; IV: intravenous; SL: sublingual

| Study | Sample size | Study description | Data report | LSD administration | LSD base doses (µg) |
| --- | --- | --- | --- | --- | --- |
| Schmid et al., 2015 [22] | N = 16 | - double-blind, placebo-controlled, randomized, crossover<br>- blood pressure, heart rate, psychomotor performance, further questionnaire assessment | 5D-ASC, 11-ASC | PO:<br>(1) 140 µg LSD base | (1) 140 |
| Carhart-Harris et al., 2016 [66] | N = 20 | - placebo-controlled, crossover, within-subject, balanced-order, double-dummy<br>- fMRI and MEG (data published in other study [77]), further questionnaire assessment | 11-ASC | IV:<br>(1) 75 µg LSD base | (1) 75 |
| Preller et al., 2017 [35] | N = 22 | - double-blind, placebo-controlled, randomized, crossover<br>- additional condition: 100 µg LSD base after 40 mg ketanserin<br>- fMRI (music paradigm) | 11-ASC | PO:<br>(1) 70 µg LSD base | (1) 70 |
| Liechti et al., 2017 [39] | N = 24 | - Healthy participants and patients with anxiety associated with life-threatening diseases | 5D-ASC, 11-ASC | PO:<br>(1) 70 µg LSD base | (1) 70 |
|  | (1) N = 16<br>(2) N = 11 (patients) | - double-blind, placebo-controlled, crossover<br>- plasma LSD levels, fMRI (resting state) (data published in [41]), further questionnaire assessment | MEQ30 | PO:<br>(1) 140 µg LSD base<br>(2) 140 µg LSD base | (1) 140<br>(2) 140 |
| Bershad et al., 2019 [44] | N = 20 | - double-blind, placebo-controlled, within-subject<br>- blood pressure, heart rate, dual n-back task, digit symbol substitution task, cyberball task, emotional images task, remote associations task, further questionnaire assessment | 11-ASC | SL:<br>(1) 6.5 µg LSD tartrate<br>(2) 13 µg LSD tartrate<br>(3) 26 µg LSD tartrate | (1) 4*<br>(2) 9*<br>(3) 18* |
| Family et al., 2020 [78] | N = 12 | - double-blind, placebo-controlled, randomized<br>- blood pressure, pulse rate, plasma LSD levels, urinalysis, ECG, balance and proprioception measures, further questionnaire assessment | 5D-ASC | PO:<br>(1) 5 µg LSD base<br>(2) 10 µg LSD base<br>(3) 20 µg LSD base | (1) 5<br>(2) 10<br>(3) 20 |
| Holze et al., 2020 [67] | N = 28 | - double-blind, placebo-controlled, crossover, double-dummy<br>- additional conditions: 125 mg MDMA, 40 mg d-amphetamine<br>- blood pressure, heart rate, plasma LSD levels, fMRI (will be published elsewhere), further questionnaire assessment | 5D-ASC, 11-ASC, MEQ30 | PO:<br>(1) 100 µg LSD base | (1) 100 |
| Hutten et al., 2020 [26] | N = 24 | - double-blind, placebo-controlled, within-subject, randomized<br>- plasma LSD levels, psychomotor vigilance task, digit symbol substitution task, cognitive control task, further questionnaire assessment | 11-ASC | PO:<br>(1) 5 µg LSD base<br>(2) 10 µg LSD base<br>(3) 20 µg LSD base | (1) 5<br>(2) 10<br>(3) 20 |
| Holze et al., 2021 [28] | N = 16 | - double-blind, placebo-controlled, crossover<br>- additional condition: 200 µg LSD base 1 h after 40 mg ketanserin<br>- blood pressure, heart rate, plasma LSD levels, further questionnaire assessment | 5D-ASC, 11-ASC, MEQ30 | PO:<br>(1) 25 µg LSD base<br>(2) 50 µg LSD base<br>(3) 100 µg LSD base<br>(4) 200 µg LSD base | (1) 25<br>(2) 50<br>(3) 100<br>(4) 200 |
| Murray et al., 2021 [43] | N = 22 | - double-blind, placebo-controlled, within-subject<br>- blood pressure, heart rate, EEG (broadband oscillatory power during resting state, event-related potentials during oddball task), further questionnaire assessment | 11-ASC | PO:<br>(1) 13 µg LSD tartrate<br>(2) 26 µg LSD tartrate | (1) 9*<br>(2) 18* |
| Wießner et al., 2021 [68] | N = 24 | - double-blind, placebo-controlled, randomized, crossover<br>- further questionnaire assessment | 11-ASC, MEQ30 | PO:<br>(1) 50 µg LSD base | (1) 50 |
| Becker et al., 2022 [79] | N = 24 | - double-blind, placebo-controlled, randomized, crossover<br>- additional condition: 40 mg ketanserin 1h after 100 µg LSD base<br>- blood pressure, heart rate, body temperature, pupil size, plasma LSD levels, plasma ketanserin levels, plasma BDNF levels, further questionnaire assessment | 5D-ASC, 11-ASC, MEQ30 | PO:<br>(1) 100 µg LSD base | (1) 100 |
| de Wit et al., 2022 [45] | N = 19 | - double-blind, placebo-controlled, randomized<br>- blood pressure, heart rate, digital symbol substitution task, *n*-back task, cyberball task, emotional images task, emotional faces task, further questionnaire assessment | 11-ASC | SL:<br>(1) 13 µg LSD tartrate<br>(2) 13 µg LSD tartrate<br>(3) 13 µg LSD tartrate<br>(4) 13 µg LSD tartrate | (1) 9*<br>(2) 9*<br>(3) 9*<br>(4) 9* |
|  | N = 19 |  |  | SL:<br>(1) 26 µg LSD tartrate<br>(2) 26 µg LSD tartrate<br>(3) 26 µg LSD tartrate<br>(4) 26 µg LSD tartrate | (1) 18*<br>(2) 18*<br>(3) 18*<br>(4) 18* |
| Family et al., 2022 [71] | (1) N = 12<br>(2) N = 7<br>(3) N = 3 | - phase 1 proof-of-concept, single-center (part 1: open-label dose-escalation study in psychedelic non-naïve participants; part 2: double-blind, placebo-controlled, randomized study in psychedelic naïve participants)<br>- additional conditions: placebo followed by 75 µg LSD base, 75 µg LSD base after 50 µg LSD base<br>- blood pressure, pulse rate, plasma LSD levels, ECG, further questionnaire assessment | 5D-ASC, MEQ30 | PO:<br>(1) 50 µg LSD base<br>(2) 75 µg LSD base<br>(3) 100 µg LSD base | (1) 50<br>(2) 75<br>(3) 100 |
| Holze et al., 2022 [5] | N = 28 | - double-blind, placebo-controlled, crossover<br>- additional conditions: 15 mg psilocybin, 30 mg psilocybin<br>- blood pressure, heart rate, plasma LSD levels, further questionnaire assessment | 5D-ASC, 11-ASC, MEQ30 | PO:<br>(1) 100 µg LSD base<br>(2) 200 µg LSD base | (1) 100<br>(2) 200 |
| Holze et al., 2023 [69] | N = 37<br>(patients) | - Patients with who experienced anxiety with or without association with a life-threatening illness<br>- 2-center, double-blind, placebo-controlled, 2-period, random-order, crossover<br>- blood pressure, heart rate, further questionnaire assessment | 5D-ASC, 11-ASC, MEQ30 | PO:<br>(1) 200 µg LSD base<br>(2) 200 µg LSD base | (1) 200<br>(2) 200 |

### Questionnaires

This meta-analysis included psychometric data from commonly applied questionnaires to assess the phenomenology of altered states of consciousness, namely from two versions of the Altered States of Consciousness Rating Scale (the 5D-ASC and the 11-ASC), and from the Mystical Experience Questionnaire (MEQ30). The Altered States of Consciousness Rating Scale [47–49] is a self-report questionnaire with 94 items rated on a visual analog scale. Two different analysis schemata are in use: In the 5D-ASC version (“5-Dimensional Altered States of Consciousness Rating Scale” [50,51]), items are assigned to five core dimensions: (1) *Auditory Alterations*, (2) *Dread of Ego Dissolution*, (3) *Oceanic Boundlessness*, (4) *Visionary Restructuralization*, and (5) *Vigilance Reduction*. In the more recent 11-ASC version (“11-factor Altered States of Consciousness Rating Scale” [52]) only 42 of the 94 questionnaires items are used in the analysis, where item scores are summarized along 11 factors: (1) *Experience of Unity*, (2) *Spiritual Experience*, (3) *Blissful State*, (4) *Insightfulness*, (5) *Disembodiment*, (6) *Impaired Control and Cognition*, (7) *Anxiety*, (8) *Complex Imagery*, (9) *Elementary Imagery*, (10) *Audio-Visual Synesthesia*, and (11) *Changed Meaning of Percepts*. Both analysis schemes have been validated and demonstrate good reliability (5D-ASC: Hoyt 0.88– 0.95 [51,53]); 11-ASC: mean Cronbach’s alpha of 0.83 [52]). The Mystical Experience Questionnaire, in its latest version the MEQ30 [54], consists of 30 items assigned to four scales: (1) *Mystical*, (2) *Positive Mood*, (3) *Transcendence of Time and Space*, and (4) *Ineffability*. This factor structure is currently recommended for analyses and has been assessed for reliability, yielding good scores for all four subscales (Cronbach’s alpha: 0.80 to 0.95) [55,56]. A more detailed description of the questionnaires can be found in Schmidt and Majić [19], Majić et al. (2015) [9] and in a recent review by de Deus Pontual et al. (2022) [21].

### Statistical analyses

We performed meta-regression analyses using restricted cubic splines. A restricted cubic spline model is non-linear and consists of a series of piecewise cubic polynomials that are connected at the position of the knots, with a linear curve before the first and after the last knot [57]. It was suggested that four or five knots are usually adequate to fully capture the underlying shape, provided the amount of included studies is sufficiently large [58,59]. Given the limited number of available studies for the 5D-ASC and MEQ30, we chose three knots for those questionnaire-data and 4 knots for the 11-ASC. To account for repeated measures and potential heteroscedasticity of error-terms resulting from repeated measurements on the same or similar set of participants, we clustered standard errors at the study level and estimated the cluster-robust variance with the small sample adjustment proposed by Pustejovsky & Tipton (2022) [60]. We assumed a within-study correlation p=0.8 for the variance-covariance matrix. The analyses were performed in R version 4.2.2 [61] with the metafor-package [62] using the restricted cubic spline model from the drc-package [63]. Cluster-robust variance was estimated with the clubSandwich-package [64] in the R robust function. The R-syntax of this analysis is available on GitHub: https://github.com/TimHirschfeld/doseresponse_LSD.

## Results

### Data description

Sixteen studies assessing the effects of LSD were included in this analysis, with overall 765 LSD applications. Table 1 contains a summary of studies included in this analysis. From the 11-ASC, 31 questionnaire datasets from 15 participant samples were included in the analysis; from the 5D-ASC, 18 datasets from nine samples were included; and from the MEQ30, 16 datasets from eight samples were included. Included studies assessed LSD doses ranging from 4 to 200 μg LSD base.

### Dose-response analyses

Radar charts for each questionnaire and dose-response relationships for each factor and scale of the respective questionnaires are presented in Figure 1 (5D-ASC and 11-ASC) and Figure 2 (MEQ30). LSD dose-dependently increased effect estimates on all factors and scales of the applied questionnaires. For data exploration and exact dose-effect-determination, the results and summary statistics of this analysis can be accessed via this interactive web-application: [Link in published article].

**Figure 1.**
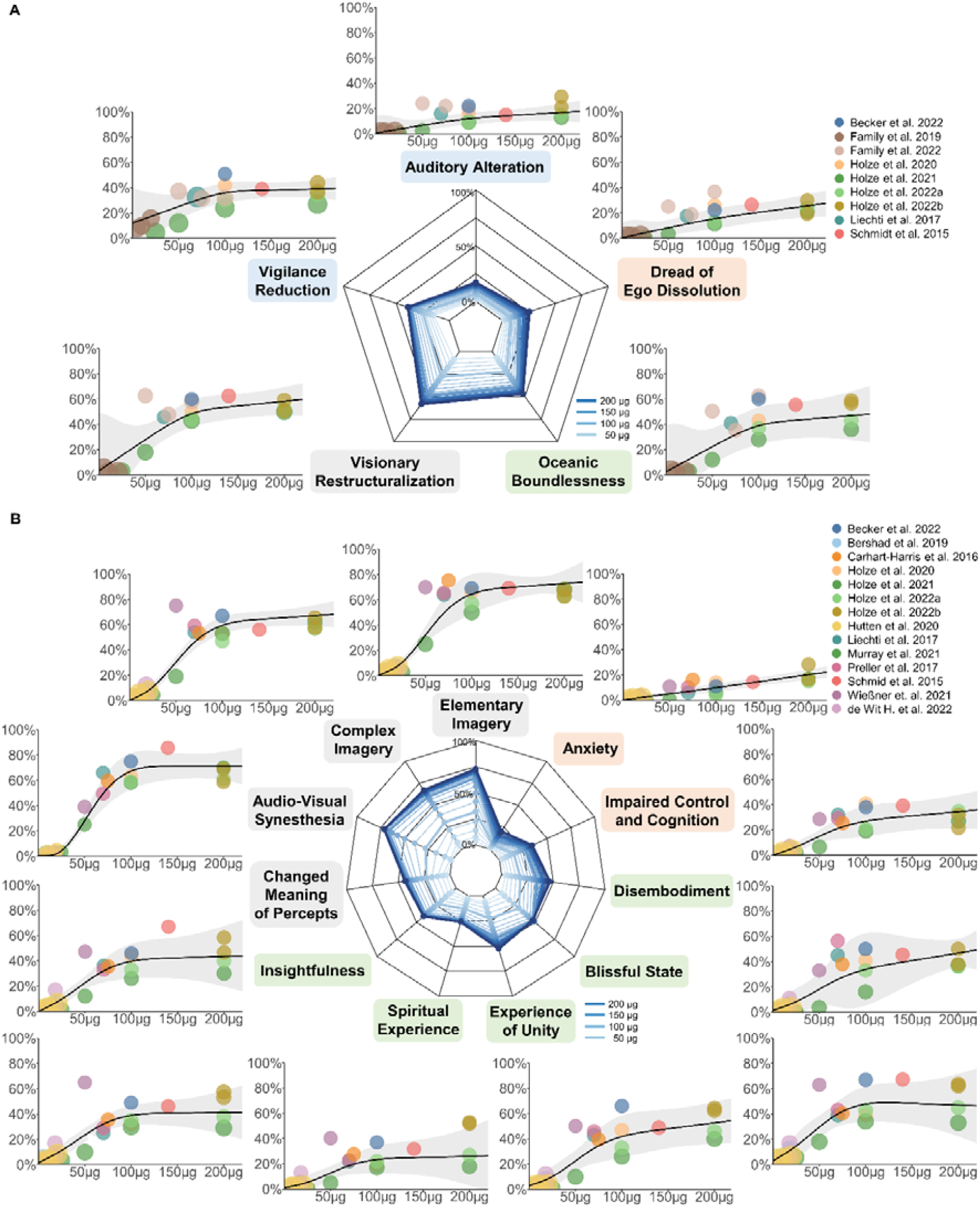
Dose-response relationships for the Altered States of Consciousness Rating Scale. **A** Dose-specific subjective effects of LSD measured with the Altered States of Consciousness Rating Scale, where questionnaire items are organized into five factors, called ‘dimensions’ of altered states of consciousness experiences (5D-ASC). **B** A finer-grained quantification of specific aspects of subjective experiences is obtained when the questionnaire is analyzed according to the 11-factors schema. These 11 factors can be considered subscales of the three core dimensions of the 5D-ASC (see corresponding colors of the subscale names). Doses are given in microgram, as absolute doses not normalized to body weight; effects are given as the percentage score of the maximum score on each factor (questionnaire items were anchored with 0% for ‘No, not more than usual’ and 100% for ‘Yes, much more than usual’). Circle color indicates from which article the data was obtained; the same color of two circles indicates statistically dependent data. Circle size corresponds to the weight of a study based on study variance (see Methods). Radar charts present the estimated dose-responses for dosages up to 200 μg. The color of individual scales corresponds to the primary dimensions and the respective subscales.

**Figure 2.**
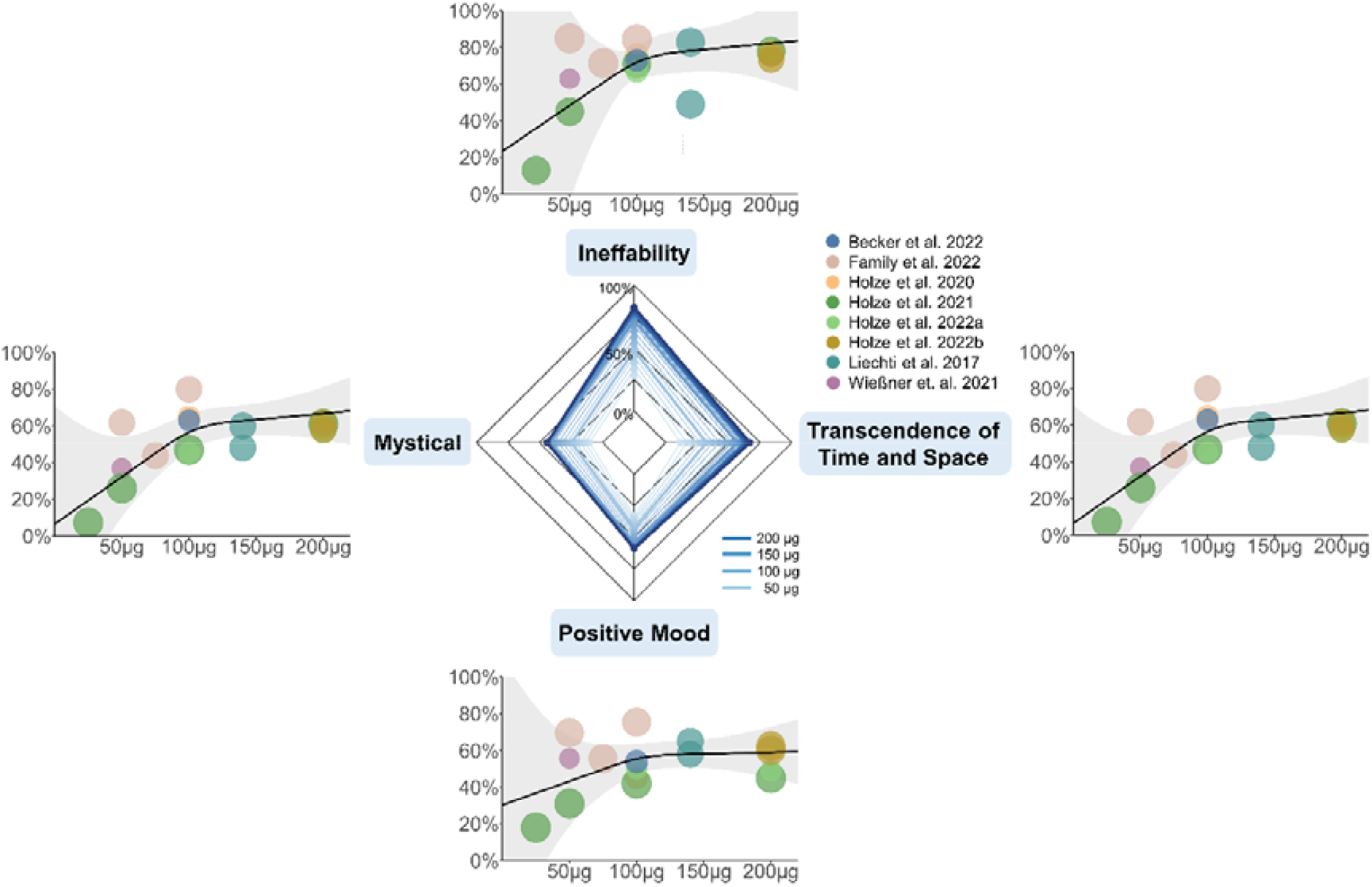
Dose-response relationships for the MEQ30. Dose-specific subjective effects of LSD measured with the Mystical Experience Questionnaire (MEQ30). Absolute doses are given in microgram. Effects on the MEQ30 are presented as the percentage score of the maximum score. Circle color indicates from which article the data was obtained; the same color of two circles indicates statistically dependent data. Circle size corresponds to the weight of the data based on study variance (see Methods). Radar charts present the estimated dose-responses for doses up to 200 μg.

## Discussion

A meta-analysis on psychometric data was performed to estimate dose-response relationships of subjective effects of LSD. The analysis included data from applications of doses from 4 to 200 μg LSD base, in healthy and clinical study participants. Given that the data is based on retrospective subjective reports relying on introspection, we did not employ assumptions on the specific shape of the fitted model (e.g., not applying a linear or a sigmoidal model); instead, we opted for fitting restricted cubic splines that allow for flexible model fits. Most of the fitted dose-response functions closely resemble sigmoid curves, which is expected and plausible for biophysiological processes. We established robust dose-response relationships, in which LSD dose increased effects on all scales and factors of the included questionnaires. On the 5D- and 11-ASC, the strongest dose-modulation was found for items referring to changes in perception and imagination (*Visionary Restructuralization, Changed Meaning of Percepts)* as well as positively experienced dissolution of boundaries between self and surroundings (*Oceanic Boundlessness*), whereas *Anxiety, Auditory Alterations* and *Dread of Ego Dissolution* were barely modulated. On the MEQ30, the strongest dose-modulation was found for *Ineffability* and the weakest for *Mystical*.

Most factors and scales exhibit a sigmoidal-like shape of dose-effects that either flattens out or reach a plateau at around 100 μg, while scales and factors with relatively small dose-modulation (*Auditory Alterations, Dread of Ego-Dissolution, Anxiety)* increase rather linearly. This is in line with a dose-response, within-subject study by Holze et al. (2021) that found similar rating scores for 100 µg and 200 µg for *Blissful State, Insightfulness*, and *Changed Meaning of Percepts* [28], but significantly higher rating scores for 200 µg compared to 100 µg on the scale *Dread of Ego Dissolution* (reflecting disembodiment and fear) [28], indicating a ceiling effect regarding positive subjective effects [65]. Despite considerable variability of effects, the present meta-analysis supports the suggestion that doses between 50 and 100 µg LSD base induce substantial psychological effects in most subjects – a dosage range currently used in most current clinical trials for therapeutic applications [65]. Accordingly, a dosage of about 100 μg LSD base most likely induces a pattern of subjective experiences that resembles what is often called a “(full) psychedelic experience”, while the likelihood of adverse reactions is kept low. Please note that the term “psychedelic experiences/state” is not used consistently and no agreement on defining criteria of a psychedelic experience have been settled on.

LSD doses below 10 μg [65] or 20 μg [24] have previously been proposed as the “microdosing” range of LSD. The fitted curves in our analysis show an onset of effects below 20 μg LSD base. For instance, visual phenomena (11-ASC: *Audio-Visual Synesthesia, Complex* and *Elementary Imagery*) show an x-axis intercept at around 10 μg, and at 20 μg reach around 10% of the maximum score. These effects are unlikely due to placebo effects, as those of the given studies which included a placebo condition reported only negligible effects [22,26,28,35,39,43– 45,66–69]. However, these estimates are strongly influenced by the fact that multiple studies investigated dosages below 20 μg, while there are no studies available in the dosage range from 20 to 50 μg, resulting in an overestimation of the curves slope at their onset. Most studies below 20 μg effectively show null-effects and make the microdosing range of 0-20 μg plausible, but more data is needed to determine the average onset of effects more precisely.

A recent systematic review indicates a mediating role of mystical-type experiences for treatment outcomes of psychedelic therapy [70], so special attention is drawn on the dose range to predict such experiences. Barrett et al. (2015) suggest that complete mystical-type experiences are reached with scores of ≥60% on each of the four factors of the MEQ30 [56]. Applying this criterion, the present analysis suggests that such experiences are rather unlikely to occur in the given dosage range. Similarly, Liechti et al. (2017) report that full mystical-type experiences occur rather rarely at LSD doses of 200 μg. The authors suggest that mystical-type and spiritual experiences might highly depend on participant characteristics and experimental setting [39], which might explain that a study by Family et al. (2022) reported of 25% and 33% of participants meeting criteria for a full mystical experience after 50 µg and 100 µg, respectively [71]. However, considering the results of the MEQ30 factors determined in this meta-analysis, datapoints of mentioned study partially lie outside or in the margins of the calculated confidence intervals and thus may not reflect general outcomes. Overall, full mystical-type experiences seem unlikely to be induced with doses below 200 µg and their occurrence appears to be strongly influenced by non-pharmacological factors. Targeted therapeutic implementation of other factors than dose need to be further investigated.

Non-pharmacological factors that influence psychedelic experiences are often categorized in set (participant personality and preparation and expectation of substance use) and setting (environment of substance administration) [30,72] and are thought to lead to considerable inter- and intra-individual variability of subjective effects [73,74]. The studies included in our analysis vary greatly with regards to experimental setting. Bershad et al. (2019) reports of “living-room style” environments with possibilities to relax, read or watch movies between measurements [44], similar to Preller et al. (2017) (“in an esthetic living-room-like room”) [35] and de Wit et al. (2022) [45]. In other studies, however, participants were also given tasks during LSD applications that involved greater effort, potentially inhibiting the manifestation of certain effects [75]. Previous work indicated that spatially constrained neuroimaging procedures may be demanding for some individuals and could increase the likelihood of challenging experiences [17]. Carhart-Harris et al. (2016) reported on fMRI and MEG measurements of over 60 min each, as well as previous MRI environment habituation and a subsequent battery of cognitive and behavioral tests [66]. From the studies included, also Holze et al. (2020) [67] and Liechti et al. (2017) [39] (fMRI data published in Müller et al., (2017) [41]) reported of fMRI measurements during the experimental session. Schmid et al. (2015) [22], Liechti et al. (2017) [39], Holze et al. (2021 [28], 2022 [5]) reported on “quiet standard hospital patient room” environments, and the study procedure by Family et al. (2022) [71] included a 60-minutes breathing exercise. Additional factors influencing the psychedelic experience and thereby increasing the variability within and between the given datasets may involve the subject’s age, previous experience with psychedelics or other mind-altering substances, as well as differences in individual pharmacokinetics [76]. The study by Family et al. (2022) for instance reports of applications of 50 µg LSD to predominantly (9 of 12) psychedelic-naïve participants, as well as 75 and 100 µg LSD to psychedelic-non-naïve participants. They report of higher questionnaire ratings on all factors of the 5D-ASC and MEQ30 in the (naïve) 50 µg cohort compared to the (non-naïve) 75 µg cohort, as well as higher ratings on most factors in the (naïve) 50 µg cohort compared to the (non-naïve) 100 µg cohort [71]. The study by Wießner et al. (2021) also reports relatively high 11-ASC scores (especially for the factors *Insightfulness, Spiritual Experience, Blissful State*, and *Complex Imaginary*) [68] compared to other studies on the same LSD dose included in this analysis. This may result from relatively high lifetime use of other psychedelics or mind-altering substances among study participants (particularly ayahuasca, with a mean lifetime use of 69 ± 131 S.D.) and the fact that most participants (67%) identify themselves as spiritual. Other studies, however, have not observed any differences in subjective effects between experienced and non-experienced LSD users when compared within the same study [5,22,28]. Therefore, differences in extra-pharmacological factors likely lead to the observed variance in the ratings of subjective experiences. The confidence intervals of the present study can yield as data-driven estimates for the expected variability of experiences, also in clinical samples (Note: The confidence intervals refer to the mean group effects, while the variance across participants can be substantially higher). Interestingly, factors assessing alterations of visual perception and cognitive abilities exhibit less variability (smaller CI range) than factors assessing experiences of unity, blissful state, and spirituality. This could be due to their highly personal character, which might be stronger influenced by non-pharmacological factors. Alternatively, variability might stem from less reliable assessment of these high-order concepts as compared to factors assessing perceptual alterations. Future studies with a more comparable assessment of non-pharmacological factors are needed to allow for systematic investigation of their exact contributions in shaping subjective experiences.

Further limitations should be considered when using the analysis at hand to determine LSD dosages. First, administration methods varied across studies. Most studies administered LSD orally as capsules, while it was administered sublingually in two studies [44,45] and intravenously in one study [66]. Moreover, LSD formulations differed across studies: most studies applied LSD base (or provided the dosage calculated for LSD base), while some dosages referred to LSD salt in the form of LSD tartrate – the typical formulation sold on the black market. No direct experimental comparison of the bioequivalence of the two formulations has been made to date. We transformed dosages according to Holze et al. (2021) [46], who suggested that 1 µg of LSD base corresponds to approximately 1.46 µg of 1:1 LSD tartrate or 1.23 - 1.33 µg 2:1 LSD tartrate, depending on the salt form and amount of crystal water [65]. Moreover, the results reflect estimates on the variability of average effects in study samples based on dosages of up to 200 µg, so the experiences of individual subjects may differ substantially. Finally, it should be noted that study samples are usually comprised of highly selected and well-prepared participants. For these reasons and because the quality and quantity of recreationally used products is often unclear, the present results do not necessarily apply to the general population or to recreational use outside controlled laboratory or therapeutic settings.

In conclusion, the present meta-analysis on subjective effects of LSD in study participants revealed dose-dependent alterations of consciousness with substantial heterogeneity of effects across laboratory and therapeutic settings on all scales and factors assessed by given questionnaires. The strongest dose-modulation was found for perceptual and imaginative changes followed by positively experienced dissolution of boundaries between self and surroundings with a limited spiritual or mystical-type character in the given dosage range. Dose-response relationships resemble sigmoidal functions on most factors and scales of the questionnaires, with an observable ceiling effect at circa 100 μg LSD base. Results may be used as a general reference to relate observed with expected dose-specific effects.

## Author contributions

TTS initiated and conceptualized the work. JP conducted the literature search and data collection. TH performed the analyses with contributions of TTS. TH, JP, TM, and TTS jointly wrote, edited, and approved the final version of the manuscript.

