## Supplementary Material for "Dose-response relationships of LSD-induced subjective experiences in humans"

for

Dose-response relationships of LSD-induced subjective experiences in humans
Hirschfeld T*, Prugger J*, Majić T, Schmidt TT
*equal contributions
Nature – Neuropsychopharmacology

**Supplementary information on the search strategy**

The meta-analysis includes psychometric questionnaire data on the subjective experience of LSD, where all data were obtained from systematic literature review procedures in line with the requirements of the Preferred Reporting Items for Systematic Reviews and Meta Analyses (PRISMA) 2020 Statement Guidelines [1]. The PRISMA flow chart in Figure S1 displays the data acquisition process. Data were included from the Altered States Database (ASDB) repository on the Open Science Framework (OSF; DOI [10.17605/OSF.IO/8MBRU](https://osf.io/8mbru/), version: "ASDB_v.2022-12-31") [2], which contains data from MEDLINE-listed studies published from 1975 until 2022-12-31. The ASDB is based on a systematic literature in line with the PRISMA statement, as reported in and described in detail in Prugger et al. (2022) [3] and Peters et al. (2023) [4]. The ASDB contains data on multiple induction methods of altered experiences, where LSD is only of those methods. Correspondingly, all data available for LSD was retrieved from the database. For this meta-analysis 28 studies were retrieved from the ASDB, of which 12 were excluded and 16 included.

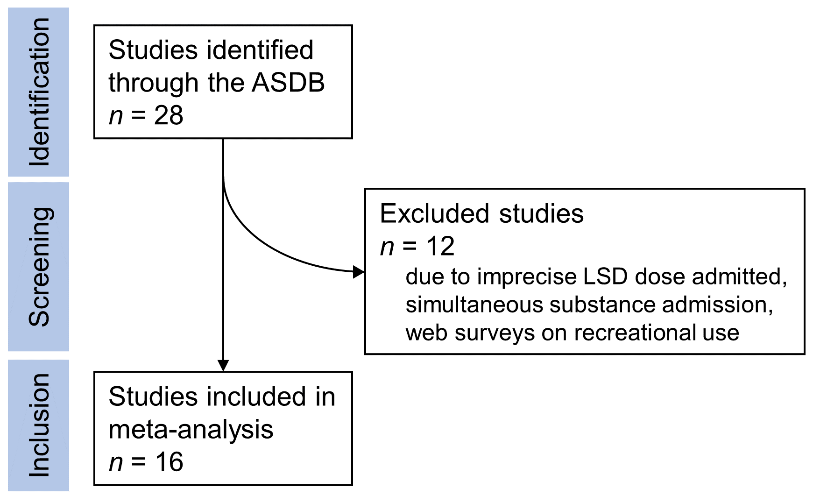
 **Supplementary Figure S1.** PRISMA 2020 Flowchart
